# Gene augmentation therapy treats mature mice with complete congenital stationary night blindness (cCSNB), improving retinal function and visual acuity

**DOI:** 10.1101/2025.07.25.666812

**Authors:** Nazarul Hasan, Cecilia A. Attaway, Mattia Di Paolo, Maureen A. McCall, Ronald G. Gregg

**Affiliations:** Departments of Biochemistry & Molecular Genetics, University of Louisville, Louisville KY 40202; Ophthalmology & Visual Science, University of Louisville, Louisville KY 40202; Anatomical Sciences & Neurobiology, University of Louisville, Louisville KY 40202

## Abstract

Recombinant adeno-associated virus (rAAV) mediated gene therapy is an effective approach for targeting therapeutic genes to retinal photoreceptors. Complete congenital stationary night blindness (cCSNB) is a genetically heterogeneous inherited retinal disease caused by mutations in one of several genes, which are part of a large and interdependent depolarizing bipolar cell (DBC) signalplex required for normal synaptic signaling with photoreceptors.

These genes include *NYX, GRM6, TRPM1, GPR179*, and *LRIT3,* and the cCSNB phenotype that results is characterized by abnormal low light vision, myopia, and nystagmus, but does not include retinal degeneration. Because of the non-progressive and recessive nature of cCSNB we investigated the potential of a gene augmentation approach in the mature retina to improve retinal function and visual acuity. We used a mouse model of cCSNB caused by the loss of LRIT3 to evaluate the efficacy of a single subretinal injection of rAAV expressing LRIT3 in either rods or cones, and the extent of restoration of retinal function and visual acuity. We show that gene augmentation by expressing LRIT3 in the rods of mature *Lrit3*^-/-^ retinas restores function and scotopic visual acuity, and when expressed on cones, improves both photopic and scotopic visual acuity.

## INTRODUCTION

Complete congenital stationary night blindness (cCSNB) is a genetically heterogeneous group of retinal disorders that cause poor night vision and are associated with high myopia and nystagmus^1^. To date, mutations in *NYX, GRM6, TRPM1, GPR179,* and *LRIT3* have been identified in humans and studied in spontaneous mutant or molecularly manipulated mice.

Mutations in each eliminate normal signaling from photoreceptors to depolarizing bipolar cells ^2^(DBCs), because they alter the expression of the postsynaptic signalplex that detects changes in glutamate release^3-13^. Most current therapeutic approaches to restore vision loss utilize rAAVs to deliver a treatment. The only clinically approved gene rAAV mediated gene augmentation therapy, Luxturna (voretigene neparvovec-rzyl), replaces expression of RPE65 to treat a form of recessive Leber’s congenital amaurosis (LCA2). Restored expression of RPE65 to the RPE, restores the visual cycle, and prevents retinal degeneration.

cCSNB is an attractive target for gene augmentation therapy because it is a recessive condition resulting from the absence of protein expression, and the retina does not degenerate, providing an intact tissue for potential therapeutic manipulation. Gene augmentation utilizing rAAVs has been used in immature retinas of some cCSNB mouse models to restore function in *Nyx, Grm6*, and *Lrit3* knockout mice ^14-19^, and in a dog *LRIT3*^-/-^ model^20^. Here we focus on LRIT3 gene augmentation in mature retina and show that treating either rods or cones with LRIT3 in mature *Lrit3*^-/-^ mice of either rods or cones improves both retina function and visual acuity.

## MATERIALS AND METHODS

### Animals

Animals were housed in the University of Louisville AALAC approved facility under a 12 hour light/12 hour dark cycle and both males and females were included in all experiments. The generation of the *Lrit3*^-/-^ mouse line has been described previously. Because there are no antibodies to Nyctalopin, staining for Nyctalopin used antibodies to GFP after crossing a transgenic line into the *Lrit3*^-/-^ background that expresses an EYFP-Nyctalopin fusion gene^21^. We refer to C57Bl6/J as Wildtype, and C57Bl6/J treated with rAAV8 RHO::GFP as Controls. For subretinal injection and ERG recording, mice were anesthetized with ketamine/xylazine solution (118/11 mg/kg, respectively) diluted in normal mouse Ringer’s solution. Mice were euthanized using CO_2_ followed by cervical dislocation, according to American Veterinary Medical Association guidelines.

### Antibodies

Antibodies used to label LRIT3, TRPM1, and mGluR6 have been described previously ^18, 22^. Others were anti-GFP (1:1000, ThermoFisher, Waltham, MA Cat# A10262), Alexa Fluor 488 Donkey anti-goat (1:1000, ThermoFisher, Waltham, MA Cat# A32814), Alexa Fluor 488 Donkey anti-rabbit (1:1000, ThermoFisher, Waltham, MA Cat# A21206), Alexa Fluor 488 Donkey anti-chicken (1:1000, Jackson ImmunoResearch Inc., West Grove PA Cat# 703545155), and Donkey anti-guinea pig Cy3 conjugated (1:1000, Millipore, Burlington, MA Cat# AP193C).

### rAAV Production and Injection

To control expression of LRIT3 using rAAVs, we used a human rhodopsin (RHO) promoter (for rods) or a mouse GNAT2 promoter (for cones), respectively ^17-19^. The expression constructs were packaged in the rAAV8 capsid by Vigene Biosciences (now Charles River Laboratories, Wilmington, MA). rAAVs (1.0µl of 1 × 10^13^ vg/ml) were introduced into mature retinas (postnatal day 35-40 (P35-P40)) using subretinal injections (www.borghuisinstruments.com). Retinal function and structure were assessed four to eight weeks after treatment.

### Electroretinography

Retinal functions were assessed by using full-field electroretinogram (ffERG) recording methods as described previously ^11, 23^. Mice were dark adapted overnight, anesthetized using ketamine/xylazine solution (118/11 mg/kg, respectively), and all procedures to prepare the mice for ERGs were performed under dim red light. Pupils were dilated by applying 0.5% phenylephrine hydrochloride or 0.1% Tropicamide solutions to the corneal surface. Body temperature was maintained by using a feedback-controlled electric heating pad (TC1000 Temperature control, CWE Inc.). Artificial tears (Tears Again, OCuSOFT, Gaithersburg, MD) were applied to the cornea, and a contact lens with a gold electrode (LKC Technologies, Gaithersburg, MD) was placed. Reference and ground needle electrodes were placed on the forehead’s midline and tail, respectively. Scotopic responses were recorded using an increasing intensity series of dim flashes (from -3.6 to 1.4 log cd s/m^2^) presented on a dark background. Photopic responses were measured by presenting a series of increasing-intensity flashes (from -0.8 to 1.4 log cd s/m^2^) after 5 minutes of light adaptation (20 cd/m^2^).

### Immunohistochemistry (IHC)

Retina wholemounts or retina transverse cryosections were prepared as described previously ^24, 25^. Briefly, mice were euthanized by CO_2_ inhalation followed by cervical dislocation, eyes were enucleated, and the cornea and lens were removed. Retinas were dissected in phosphate buffered saline (PBS, pH 7.4) and fixed for 15-60 min in 4% paraformaldehyde diluted in 0.1M PB, washed 3X with PBS 10 min each, and cryoprotected in a graded series of sucrose solutions (5, 10, 15 and 20% in PBS) and finally in OCT:20% sucrose (2:1).

Cryoprotected retinas were frozen and transverse retinal sections (18μm) were cut on a cryostat (Leica Biosystems, Buffalo Grove, IL), mounted on Superfrost Plus slides (Thermo Fisher Scientific, Waltham, MA), and stored at -80°C. Immunohistochemistry methods have been described previously ^24^. Retina sections were imaged on an Olympus FV-4000 Confocal Microscope and contrast and brightness adjusted using Fluoview Software (Olympus, Waltham, MA).

### Visual Acuity

Visual acuity was measured using a two-alternative forced choice method ^26^ modified also to measure scotopic visual acuity ^27^. Mice are placed in shallow water in a trapezoidal tank displaying a grating on one of 2 computer monitors and trained to choose the screen with the grating at a fixed distance to escape to a submerged platform. We train mice under photopic conditions (screens 100 cd/m^2^), then dark adapt them for 1 hour at 0.01 cd/m^2^, and repeat the task with both screens at 0.025 cd/m^2^, obtaining the scotopic visual acuity.

### Quantification and Statistical Analysis Rod counts

The percentage of rod LRIT3 expressing synapses in rAAV8 RHO::Lrit3 treated *Lrit3*^-/-^ retinas was determined from whole mount retinas after staining for mGluR6 and LRIT3. Confocal images using a 40x (NA = 1.4) objective on an Olympus FV4000 were collected from 5 areas (central and 4 quadrants) in each retina and averaged. Counts were made from maximum projections of z-stacks of whole mount images using ImageJ. The total number of rod synapses (mGlur6 positive) and those positive for LRIT3 were counted.

### Statistical Analysis

Prism 10.5.0 (774) (GraphPad Software, Inc., La Jolla, CA) was used for statistical analyses (see text and figure legends). Šídák’s multiple comparison test was used to adjust for multiple testing when appropriate, and the p_adj_ is reported. Statistical significance, p_adj_≤ 0.05.

## RESULTS

To express LRIT3 in rods or cones, we constructed rAAV vectors (Fig. 1A) that use either a human rhodopsin (RHO) ^28^ or a GNAT2 ^29^ promoter, respectively. In this study, we evaluated if rod-or cone-mediated retinal function and visual acuity could be restored by expressing LRIT3 in mature *Lrit3*^-/-^ mice at P35 after the visual critical period ^30^.

**Figure 1.**
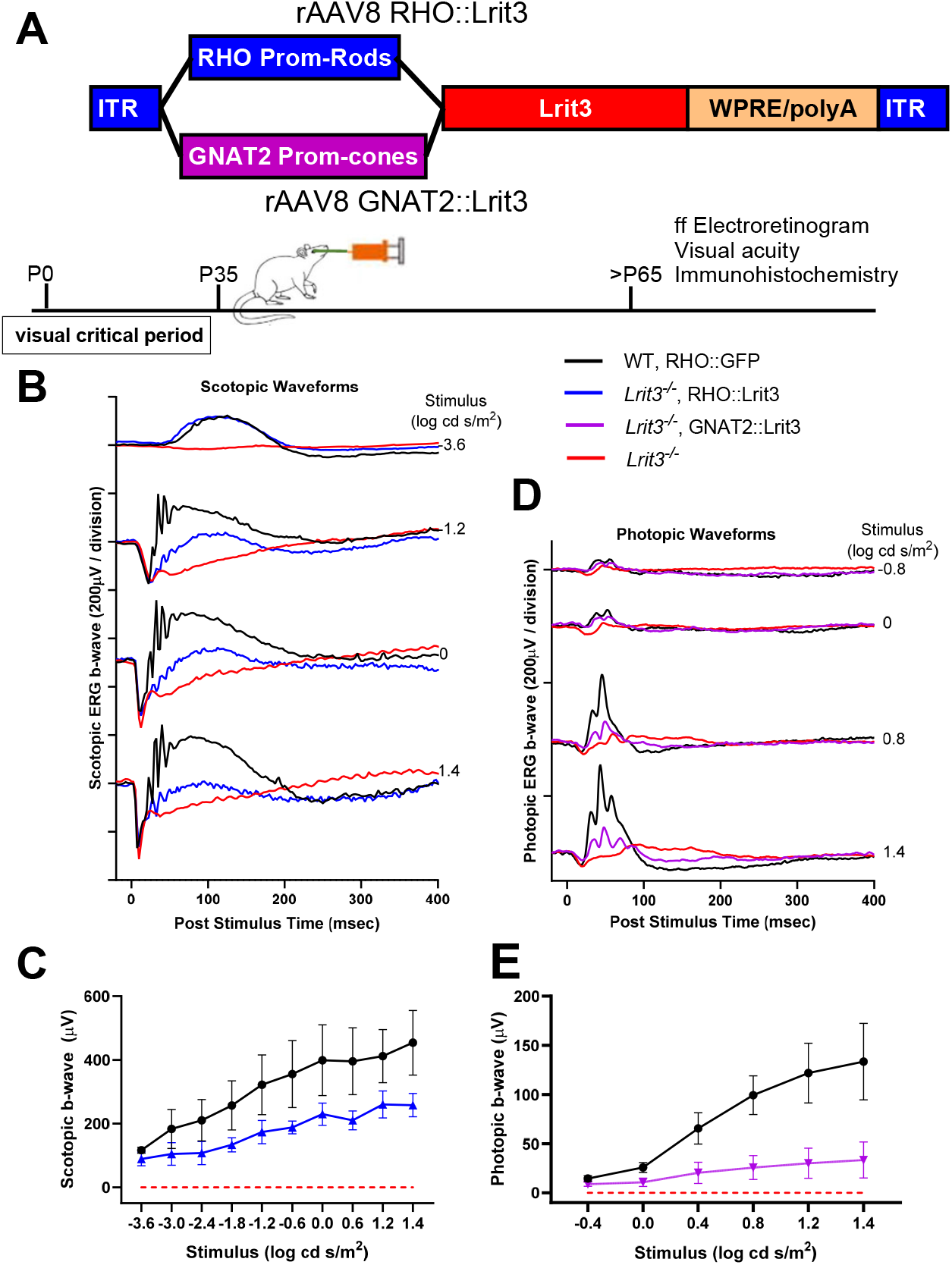
LRIT3 expression in rods or cones restores retinal function. **A.** Schematic of the two rAAV vectors and experimental design used in the study. **(B)** Scotopic ffERG waveforms of typical Control (WT, RHO::GFP, n=8); *Lrit3*^-/-^ treated (*Lrit3*^-/-^, RHO::Lrit3, n=13); untreated *Lrit3*^-/-^, n=6), and **(C)** summary data of the amplitudes of ffERG under scotopic conditions of all mice.**(D)** Photopic ffERG waveforms of typical responses from single Control (WT, RHO::GFP, n=8) *Lrit3*^-/-^ treated (*Lrit3*^-/-^, GNAT2::Lrit3, n=20) and untreated *Lrit3*^-/-^, n= 6) mouse at several flash intensities, and **(E)** summary data of the amplitudes of ffERG under photopic conditions of all mice. Values are mean±SD

### LRIT3 expression in mature rod photoreceptors of *Lrit3*^-/-^ restores rod retinal function

Four to six weeks after subretinal injection of rAAV8 RHO::Lrit3 in mature *Lrit3*^-/-^ mice, we measured scotopic and photopic full-field electroretinograms (ffERGs) to assess rod and cone driven function, respectively. Compared to *Lrit3*^-/-^ mice, *Lrit3*^-/-^ mice treated with RHO::Lrit3 had increased rod-isolated ffERG b-wave amplitudes as well as at all other tested scotopic flash intensities (Fig. 1B,C; p_adj_<0.001). Compared to Control the b-wave amplitudes remained significantly smaller (WT treated with rAAV8 RHO::GFP ; p_adj_<0.001 for all flash intensities; Fig. 1B, C). As expected, rAAV8 RHO::Lrit3 treatment of *Lrit3*^-/-^ retinas did not restore a photopic b-wave, which remained similar to that in *Lrit3*^-/-^ mice (p_adj_>0.05 for all flash intensities; see Supplementary Fig. 1 for data). This reinforces the specificity of our RHO promoter, which restricts expression to rods, as reported previously ^19^.

*Lrit3*^-/-^ mice treated with GNAT2::Lrit3 had increased photopic ffERG b-wave amplitudes at all tested flash intensities (Fig. 1D,E; p_adj_<0.001 all other flash intensities). Consistent with our previous report^16^, response amplitudes remained significantly below Controls (p_adj_<0.001 for all flash intensities).

### LRIT3 expression in mature *Lrit3*^-/-^ rod photoreceptors restores expression of TRPM1 and Nyctalopin to postsynaptic rod BC cell dendrites

Treatment with rAAV8 RHO::Lrit3 restored DBC signalplex postsynaptic protein localization consistent with the functional ERG rescue. Both Nyctalopin and TRPM1 expression were restored to rod BC dendrites (Figure 2A, B), and co-localized with the rescued expression of LRIT3 puncta in the OPL, a pattern similar to Wildtype (Fig. 2Aiii and 2Biii).

**Figure 2.**
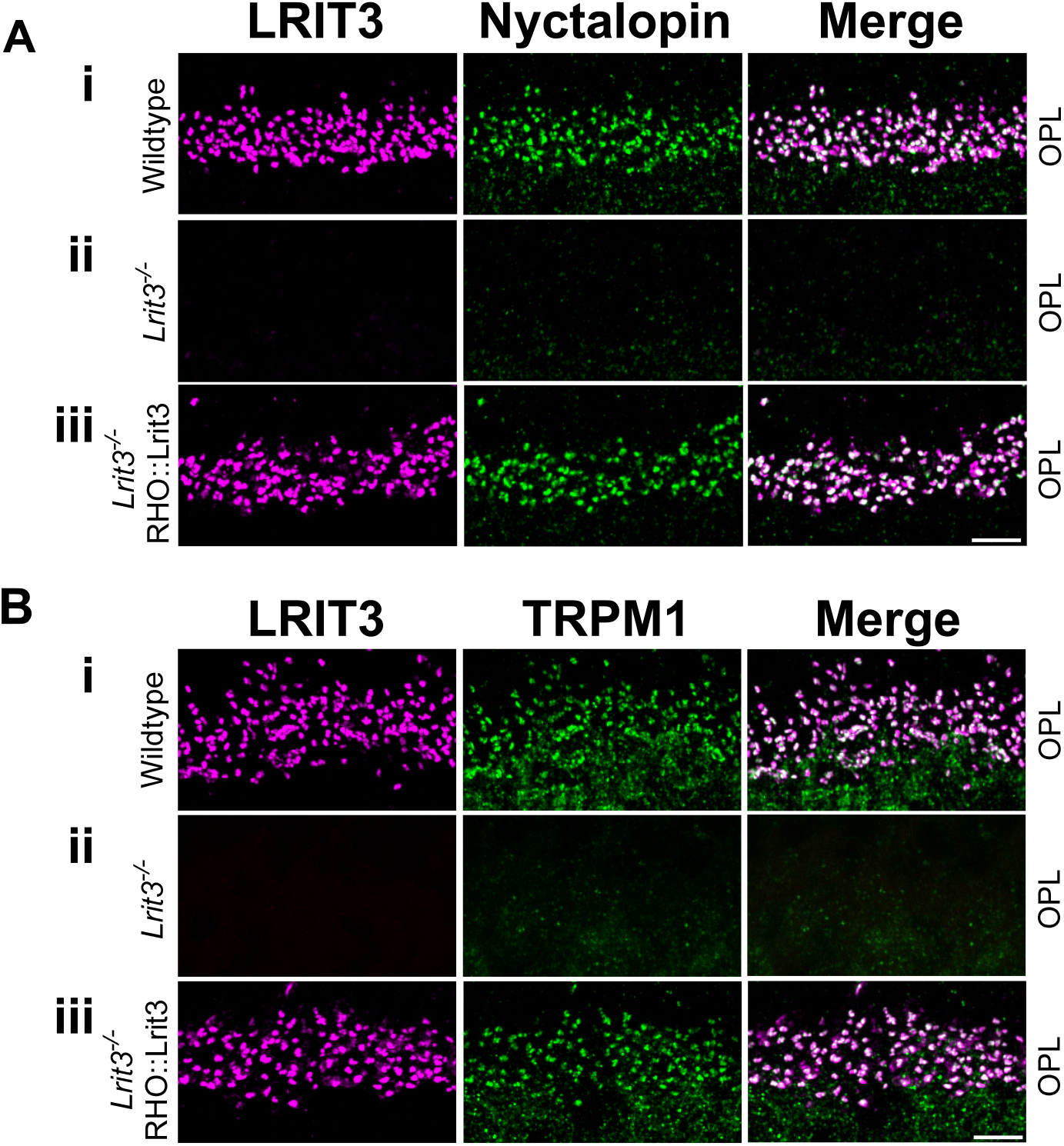
LRIT3 expression in the rods of *Lrit3*^-/-^ mice restores DBC signalplex components. Representative images of the outer plexiform layer (OPL) after immunohistochemistry of transverse retinal sections from (i) Wildtype, (ii) *Lrit3*^-/-^ and, (iii) *Lrit3*^-/-^ retinas treated with rAAV8 RHO::LRIT3 using antibodies to **(A)** LRIT3 (magenta), GFP-Nyctalopin (green) and merge; and **(B)** LRIT3 (magenta), TRPM1 (green) and merge. Scale bar = 5µm. Images are representative of four retinas.

To determine if there is a relationship between the scotopic ERG b-wave amplitude and the number of rods whose function is rescued, we quantified rAAV infection frequency in retinal wholemounts by staining for mGluR6 and LRIT3 expression (Fig. 3). In Wildtype retinas, mGluR6 and LRIT3 colocalize (Fig 3Ai), whereas in *Lrit3*^-/-^ rod-rod BC synapses only mGluR6 is present (Fig 3Aii). In retinas treated with rAAV8 RHO::Lrit3, infected rods show puncta double-labeled with mGluR6 and LRIT3, whereas uninfected rod puncta show only staining by mGluR6 (Fig. 3iii, 4 examples within white circles). Quantification of these data from eight treated retinas shows that the infection percentage across RHO::Lrit3 treated retinas ranged from 22-47% (Fig. 3B). To determine if there was a correlation between structure and function, we plotted the percentage of rods expressing LRIT3 against the dark-adapted, rod isolated ffERG (flash intensity -3.6 log cd.s/m^2^, Fig. 3B) for treated and untreated *Lrit3*^-/-^ and Widtype retinas (Fig. 3B). What is notable is that treatment rescued the ffERG b-wave in all treated *Lrit3*^-/-^ retinas, and the amplitudes fall above the line connecting the *Lrit3*^-/-^ and Wildtype retinas (Fig. 3B), suggesting that the relationship between the ffERG b-wave amplitude and the number of infected rods expressing LRIT3 is not linear.

**Figure 3.**
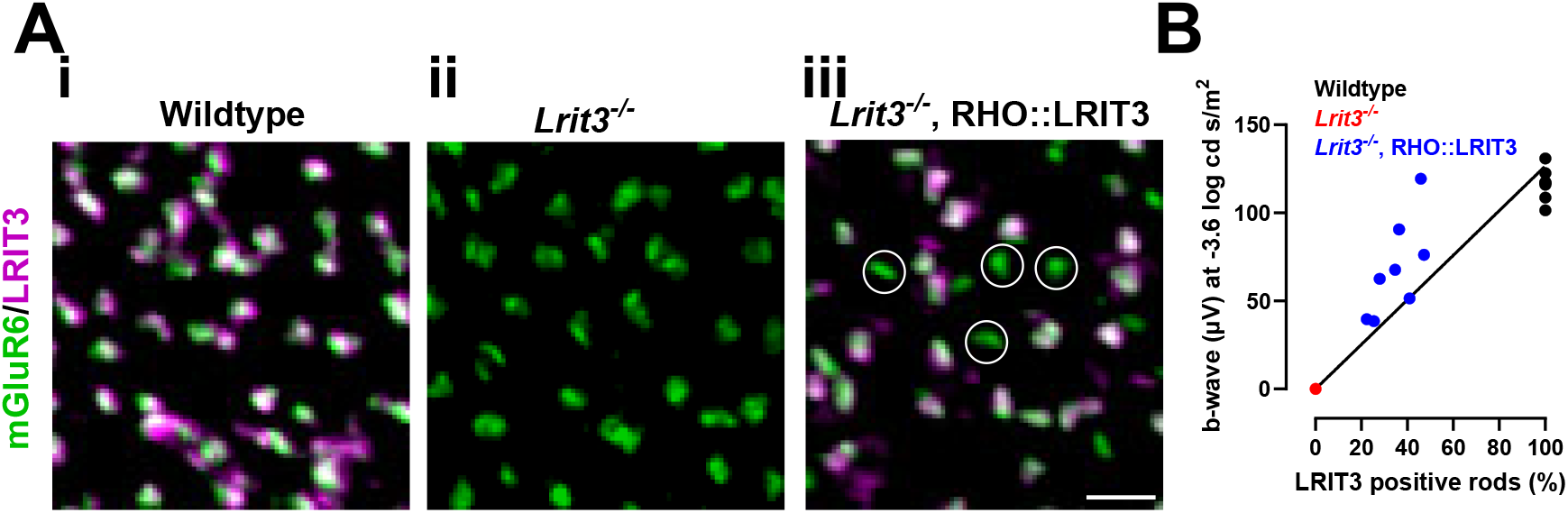
The fraction of rods with restored LRIT3 expression correlates with the amplitude of the scotopic ffERG b-wave. **(A)** Representative image of wholemount retina taken at the level of the OPL from Wildtype, *Lrit3^-/-^, Lrit3*^-/-^ treated with rAAV8 RHO::Lrit3, and stained with antibodies to LRIT3 (magenta) and mGluR6 (green). White circles in the treated retina represent rod spherules that do not express LRIT3. Scale bar = 5µm. **(B)**. Amplitude of rod-isolated ffERG b-wave versus percent of rods expressing LRIT3 (n=8). Line is between *Lrit3*^-/-^ and Wildtype. Scale bar = 5µm. Images are representative of eight retinas.

### Visual acuity in *Lrit3*^-/-^ mice improves after rescue of rods or cones

Similar to these results showing restoration of function in mature LRIT3 rods, we previously showed that photopic ERG function and cone signalplex protein expression are rescued when LRIT3 is expressed in cones of mature *Lrit3*^-/-^ mice^16^. A key question in any therapy that treats the mature retina is whether it is sufficiently early (within the critical period) to improve overall vision. To address this question, we used a two-alternative forced-choice visual discrimination task to measure visual acuity mediated by rescued *Lrit3*^-/-^ cones ^26^, and we adapted this test to evaluate visual acuity mediated by rescued rods.

The mice were trained, and visual acuity was measured under photopic, rod-saturating conditions, followed by measurements under scotopic conditions, 60 minutes of dark adaptation (Fig. 4, white and grey bars represent photopic and scotopic visual acuity, respectively). The treated mice were compared to Wildtype and untreated *Lrit3*^-/-^ age-matched mice (all p-values for all comparisons in this dataset are in Supplementary Table 1). Wildtype mice have similar visual acuity under both photopic and scotopic conditions (p_adj_=0.416).

**Figure 4.**
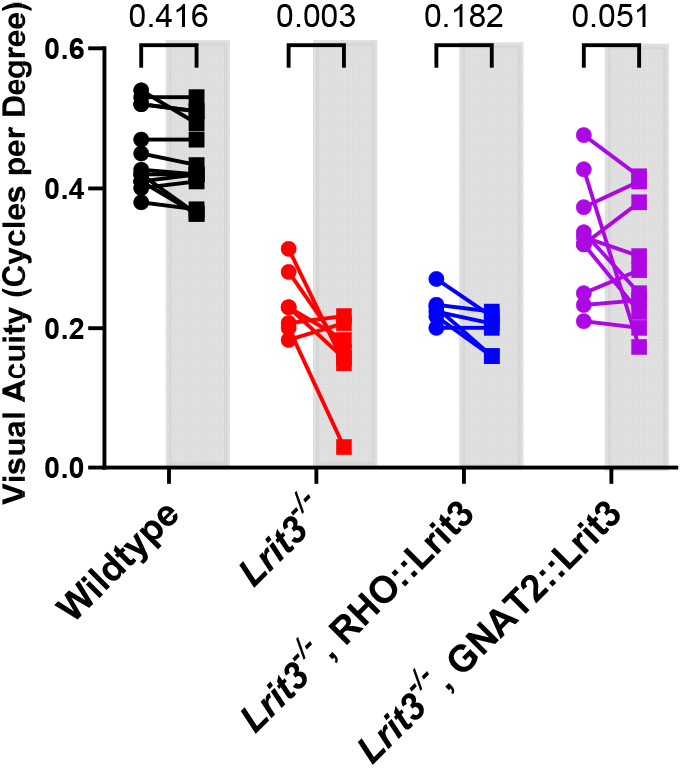
Expression of LRIT3 in rods or cones of *Lrit3*^-/-^ mice improves visual acuity under scotopic conditions. Wildtype (n=13), *Lrit3*^-/-^ (n=6), rAAV RHO::Lrit3 treated *Lrit3*^-/-^ (n=6) and rAAV8 GNAT::LRIT3 treated *Lrit3*^-/-^ (n=10). The white and gray bars indicate photopic and scotopic visual acuity, resepctively. P values are p_adj_ values from two-way ANOVA after adjustment with Šídák’s multiple comparisons test (Full statistics Supplemental Table 1).

Untreated *Lrit3*^-/-^ mice have significantly poorer photopic and scotopic visual acuity compared to Wildtype (p_adj_ values both <0.001), and their scotopic visual acuity is poorer still compared to their photopic visual acuity (p_adj_=0.003). Treatment with rAAV8 RHO::Lrit3 restored scotopic visual acuity to photopic levels with rAAV8 RHO::Lrit3 (p_adj_ = 0.182). In contrast, treatment with rAAV8 GNAT2::Lrit3 improved both photopic and scotopic visual acuity, compared to untreated *Lrit3*^-/-^ mice (photopic: p_adj_<0.023; scotopic p_adj_<0.001). Notably, three rAAV8 GNAT2::Lrit3 treated *Lrit3*^-/-^ mice had visual acuities similar to Wildtype.

In summary, these data show that AAV mediated expression of LRIT3 in either rods or cones, restores the post-synaptic signaling complex on DBCs and this translates to improved ffERG b-waves, and dark adapted visual acuity. When cones are treated, both photopic and scotopic visual acuity are restored to near wildtype acuity in many animals.

## DISCUSSION

Gene therapy for IRDs (inherited retinal diseases) is a proven approach, with FDA approval in 2017 of Luxturna, a rAAV approach to treat mature and juvenile patients with a form of Leber’s congenital amaurosis (LCA2), caused by biallelic null mutations in *RPE65* and better outcomes were reported when younger patients are treated ^31^. This indicates that the developmental stage of the retina may predict the outcome, however, this interpretation is clouded by differences in the stage of photoreceptor degeneration. In contrast, cCSNB is a stationary disease characterized by poor night vision from birth. The recessive and non-progressive nature of cCSNB allows us to interpret our rescue results as a function of retinal maturity alone.

Normal vision requires photoreceptors to communicate with inner retinal neurons by altering glutamate release from their axon terminals. The changes in glutamate release are detected by the DBCs as well as the hyperpolarizing BCs (HBCs) that form the ON and OFF retinal pathways, respectively. However, there is crosstalk between the ON and OFF pathways, and this is reflected by defects in ON pathway signaling that result in defects in the OFF pathway in mouse models of cCSNB^18^.

AAV mediated gene augmentation in mouse models of cCSNB caused by mutations in *Grm6, Nyx*, and *Lrit3* have produced varying degrees of successful rescue. Some of the variability in the results can be attributed to the size of the gene (e.g., Grm6 is large and difficult to package efficiently) and to the target cells (e.g., DBCs are more difficult to infect). Other variation results from differing treatment ages. For example, when DBCs in *Grm6* and *Nyx* mutants were delivered in young mice, P15 and P5, respectively ^14, 15^, only a small improvement in the scotopic ERG b-wave amplitude was observed. Results for LRIT3 mediated rescue in cCSNB, show differing results dependent on the study. One, delivered AAVs designed to express LRIT3 at P30 to DBCs, photoreceptors, or both by intravitreal injections, and only a small improvement in the scotopic ffERG b-wave, and no improvement in the photopic ffERG b-wave was observed. Despite this result, the mice had improved optomotor responses, suggesting improvements in subcortical vision^32^. In contrast, a second study using earlier intervention (P5), expressing LRIT3 in rods of *Lrit3*^-/-^ mice improved the scotopic ffERG b-wave, although not to that of controls ^18^. Furthermore, late-intervention treatment of cones in mature mouse retinas markedly improved the photopic ffERG b-wave, although the impact on vision was not reported.^17^ A study using an LRIT3 mutant dog model of cCSNB delivered an rAAV optimized to infect dog DBCs between 1.3 and 2.8 years of age, resulting in an improvement in the scotopic ffERG b-wave and a rod-mediated visual mobility task ^20^. This study was done prior to the demonstration that LRIT3 is expressed in photoreceptors^19^. Taken together, these studies suggest that gene augmentation for LRIT3-mediated cCSNB can improve some functional measures; however, they do not systematically evaluate the impact of these interventions in the mature retina, and none have evaluated cortically mediated visual acuity. This prompted us to evaluate the outcome of targeted gene augmentation treatment to mature rods and cones after the visual critical period and assess the impact on visual acuity.

The improvement in photopic visual acuity when LRIT3 is expressed in cones of *Lrit3*^-/-^ mice is not that surprising, but their improvement under scotopic conditions was unexpected. This improvement in scotopic acuity was not caused by the small number of rods (∼7%) also rescued due to promoter leakiness, because even when rod-specific promoters are used and 50% of rod signaling is restored, scotopic acuity remains like that of untreated *Lrit3*^-/-^ animals (Fig. 4).

Furthermore, these data suggest that the treatment of cones provides some animals with near Wildtype visual acuity, despite a lack of signaling through rod BCs. Given that the number of cones expressing LRIT3 is similar after treatment, further experiments are needed to understand the details of this variability.

While the primary phenotype of cCSNB is poor scotopic vision, many patients and cCSNB mouse models have nystagmus^33^. In the mouse models, this has been shown to result from abnormal retinal activity, which manifests as retinal oscillations, first described in the *Nyx*^*nob*^ mouse ^34, 35^ and now demonstrated in LRIT3 and mGluR6 deficient lines ^36^. We also found that this oscillation was decreased in BCs cells from *Lrit3*^-/-^ mice after expressing LRIT3 in some rods ^18^. Therefore, it is possible that the visual acuity improvements we document here could be a combination of restoring normal signaling between photoreceptors and DBCs, and a decrease in the abnormal retinal activity that may be driving nystagmus.

The improvement in both scotopic and photopic visual acuity demonstrated in this mouse model of cCSNB sets the stage for additional studies on the ability to rescue other cCSNB models, and a related disorder, incomplete CSNB (iCSNB), which is also caused by a disruption in rod and cone synaptic function ^37^. As gene therapy for IRDs progresses, both complete and incomplete CSNB patients may be treated, improving scotopic and photopic visual acuity and decreasing nystagmus. Further, these treatments are likely to be effective in adult patients with cCSNB.

## DATA AVAILABILITY

Data generated and analyzed during this study can be found within the published article. Raw data and images can be provided by the corresponding author upon reasonable request.

## ACKNOWLEDGEMENTS

We want to thank Timothy Hoffman for his technical assistance.

## AUTHOR CONTRIBUTIONS

Conceptualization, RGG and NH; Investigation, NH.; CA.; MD. ; Writing, NH, RGG, MAM.

## ETHICS APPROVAL

All procedures were performed in accordance with the Society for Neuroscience policies on the use of animals in research and the University of Louisville Institutional Animal Care and Use Committee.

## REFERENCES

1. Zeitz C, Robson AG, Audo I. Congenital stationary night blindness: an analysis and update of genotype-phenotype correlations and pathogenic mechanisms. Progress in retinal and eye research 2015; 45: 58–110.

2. Ray TA. Ph.D. Thesis. Constructing the rod bipolar signalplex using animal models of retinal dysfunction. https://ir.library.louisville.edu/etd/1188/. 2013.

3. Masu M, Iwakabe H, Tagawa Y, Miyoshi T, Yamashita M, Fukuda Y et al. Specific deficit of the ON response in visual transmission by targeted disruption of the mGluR6 gene. Cell 1995; 80(5):757–65.

4. Pusch CM, Zeitz C, Brandau O, Pesch K, Achatz H, Feil S et al. The complete form of X-linked congenital stationary night blindness is caused by mutations in a gene encoding a leucine-rich repeat protein. Nat Genet 2000; 26(3): 324–7.

5. Bech-Hansen NT, Naylor MJ, Maybaum TA, Sparkes RL, Koop B, Birch DG et al. Mutations in NYX, encoding the leucine-rich proteoglycan nyctalopin, cause X-linked complete congenital stationary night blindness. Nat Genet 2000; 26(3): 319–23.

6. Gregg R, Lukasiewicz P, Peachey N, Sagdullaev B, McCall M. Nyctalopin is required for signaling through depolarizing bipolar cells in the murine retina. Investigative Ophtalmology and Visual Science 2003; 44(5): 4180.

7. Morgans CW, Zhang J, Jeffrey BG, Nelson SM, Burke NS, Duvoisin RM et al. TRPM1 is required for the depolarizing light response in retinal ON-bipolar cells. Proc Natl Acad Sci U S A 2009; 106(45): 19174–8.

8. Shen Y, Heimel JA, Kamermans M, Peachey NS, Gregg RG, Nawy S. A transient receptor potential-like channel mediates synaptic transmission in rod bipolar cells. J Neurosci 2009; 29(19): 6088–93.

9. Nakamura M, Sanuki R, Yasuma TR, Onishi A, Nishiguchi KM, Koike C et al. TRPM1 mutations are associated with the complete form of congenital stationary night blindness. Mol Vis 2010; 16: 425–37.

10. Audo I, Bujakowska K, Orhan E, Poloschek CM, Defoort-Dhellemmes S, Drumare I et al. Whole-Exome Sequencing Identifies Mutations in GPR179 Leading to Autosomal-Recessive Complete Congenital Stationary Night Blindness. Am J Hum Genet 2012; 90(2): 321–30.

11. Peachey NS, Ray TA, Florijn R, Rowe LB, Sjoerdsma T, Contreras-Alcantara S et al. GPR179 is required for depolarizing bipolar cell function and is mutated in autosomal-recessive complete congenital stationary night blindness. Am J Hum Genet 2012; 90(2): 331–9.

12. Zeitz C, Jacobson SG, Hamel CP, Bujakowska K, Neuille M, Orhan E et al. Whole-exome sequencing identifies LRIT3 mutations as a cause of autosomal-recessive complete congenital stationary night blindness. Am J Hum Genet 2013; 92(1): 67–75.

13. Neuille M, El Shamieh S, Orhan E, Michiels C, Antonio A, Lancelot ME et al. Lrit3 deficient mouse (nob6): a novel model of complete congenital stationary night blindness (cCSNB). PLoS One 2014; 9(3): e90342.

14. Scalabrino ML, Boye SL, Fransen KM, Noel JM, Dyka FM, Min SH et al. Intravitreal delivery of a novel AAV vector targets ON bipolar cells and restores visual function in a mouse model of complete congenital stationary night blindness. Hum Mol Genet 2015; 24(21): 6229–39.

15. Varin J, Bouzidi N, Dias MMS, Pugliese T, Michiels C, Robert C et al. Restoration of mGluR6 Localization Following AAV-Mediated Delivery in a Mouse Model of Congenital Stationary Night Blindness. Invest Ophthalmol Vis Sci 2021; 62(3): 24.

16. Hasan N, Gregg RG. Cone Synaptic function is modulated by the leucine rich repeat (LRR) adhesion molecule LRFN2. eNeuro 2024; 11(3).

17. Gregg RG, Hasan N, Borghuis BG. LRIT3 expression in cone photoreceptors restores post-synaptic bipolar cell signalplex assembly and partial function in Lrit3 (-/-) mice. iScience 2023; 26(4): 106499.

18. Hasan N, Pangeni G, Ray TA, Fransen KM, Noel J, Borghuis BG et al. LRIT3 is Required for Nyctalopin Expression and Normal ON and OFF Pathway Signaling in the Retina. eNeuro 2020; 7(1).

19. Hasan N, Pangeni G, Cobb CA, Ray TA, Nettesheim ER, Ertel KJ et al. Presynaptic Expression of LRIT3 Transsynaptically Organizes the Postsynaptic Glutamate Signaling Complex Containing TRPM1. Cell Rep 2019; 27(11): 3107–3116 e3.

20. Miyadera K, Santana E, Roszak K, Iffrig S, Visel M, Iwabe S et al. Targeting ON-bipolar cells by AAV gene therapy stably reverses LRIT3-congenital stationary night blindness. Proc Natl Acad Sci U S A 2022; 119(13): e2117038119.

21. Gregg RG, Kamermans M, Klooster J, Lukasiewicz PD, Peachey NS, Vessey KA et al. Nyctalopin expression in retinal bipolar cells restores visual function in a mouse model of complete X-linked congenital stationary night blindness. J Neurophysiol 2007; 98(5): 3023–33.

22. Ronald G. Gregg NH, Bart G. Borghuis. LRIT3 expression in cone photoreceptors restores post-synaptic bipolar cell signalplex assembly and function in Lrit3-/-mice. iScience 2023.

23. Ray TA, Heath KM, Hasan N, Noel JM, Samuels IS, Martemyanov KA et al. GPR179 is required for high sensitivity of the mGluR6 signaling cascade in depolarizing bipolar cells. J Neurosci 2014; 34(18): 6334–43.

24. Hasan N, Pangeni G, Cobb CA, Ray TA, Nettesheim ER, Ertel KJ et al. Presynaptic Expression of LRIT3 Transsynaptically Organizes the Postsynaptic Glutamate Signaling Complex Containing TRPM1. Cell Rep 2019; 27(11): 3107–3116.e3.

25. Hasan N, Ray TA, Gregg RG. CACNA1S expression in mouse retina: Novel isoforms and antibody cross-reactivity with GPR179. Visual Neurosci 2016; 33.

26. Prusky GT, West PW, Douglas RM. Behavioral assessment of visual acuity in mice and rats. Vision Res 2000; 40(16): 2201–9.

27. Attaway CA. Advancing research on Retinitis Pigmentosa: An investigation of natural history, visual system plasticity, and mutation-independent treatment. Ph.D., University of Louisville, https://ir.library.louisville.edu/etd/4564, 2025.

28. Mussolino C, della Corte M, Rossi S, Viola F, Di Vicino U, Marrocco E et al. AAV-mediated photoreceptor transduction of the pig cone-enriched retina. Gene Ther 2011; 18(7): 637–45.

29. Drinnenberg A, Franke F, Morikawa RK, Juttner J, Hillier D, Hantz P et al. How Diverse Retinal Functions Arise from Feedback at the First Visual Synapse. Neuron 2018; 99(1): 117–134 e11.

30. Hooks BM, Chen C. Critical periods in the visual system: changing views for a model of experience-dependent plasticity. Neuron 2007; 56(2): 312–26.

31. Fischer MD, Simonelli F, Sahni J, Holz FG, Maier R, Fasser C et al. Real-World Safety and Effectiveness of Voretigene Neparvovec: Results up to 2 Years from the Prospective, Registry-Based PERCEIVE Study. Biomolecules 2024; 14(1).

32. Varin J, Bouzidi N, Gauvain G, Joffrois C, Desrosiers M, Robert C et al. Substantial restoration of night vision in adult mice with congenital stationary night blindness. Mol Ther Methods Clin Dev 2021; 22: 15–25.

33. Kamermans M, Winkelman BHJ, Holzel MB, Howlett MHC, Kamermans W, Simonsz HJ et al. A retinal origin of nystagmus-a perspective. Front Ophthalmol (Lausanne) 2023; 3: 1186280.

34. Demas J, Sagdullaev BT, Green E, Jaubert-Miazza L, McCall MA, Gregg RG et al. Failure to maintain eye-specific segregation in nob, a mutant with abnormally patterned retinal activity. Neuron 2006; 50(2): 247–59.

35. Winkelman BHJ, Howlett MHC, Holzel MB, Joling C, Fransen KH, Pangeni G et al. Nystagmus in patients with congenital stationary night blindness (CSNB) originates from synchronously firing retinal ganglion cells. PLoS Biol 2019; 17(9): e3000174.

36. Holzel MB, Kamermans W, Winkelman BHJ, Howlett MHC, De Zeeuw CI, Kamermans M. A common cause for nystagmus in different congenital stationary night blindness mouse models. J Physiol 2023; 601(23): 5317–5340.

37. Zhang Y, Lin S, Yu L, Lin X, Qu S, Ye Q et al. Gene therapy shines light on congenital stationary night blindness for future cures. J Transl Med 2025; 23(1): 392.

