## Supplementary material for "Gene augmentation therapy treats mature mice with complete congenital stationary night blindness (cCSNB), improving retinal function and visual acuity": Supp Table 1 and Supp Figure 1

### Supplementary Figure 1

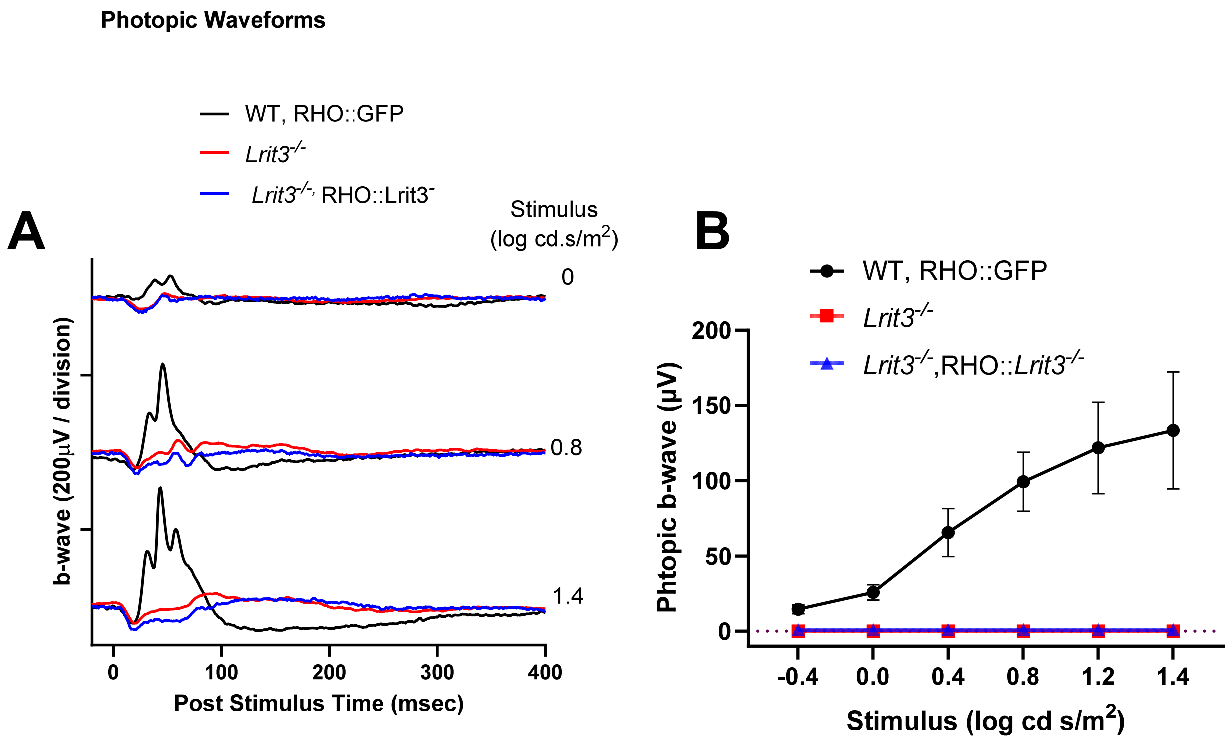

**Supplementary Figure 1.** **(A)** Photopic ffERG waveforms for an example of each ex[eriemntal group. Control; *Lrit3*$^{-/-}$ ; rAAV8 RHO::Lrit3 treated *Lrit3*$^{-/-}$ mouse. **(B)** Summary data for photopic fERG for all mice, Control (n=8); *Lrit3*$^{-/-}$ (n=6); rAAV8 RHO::Lrit3 treated *Lrit3*$^{-/-}$ mouse Note that the treatment does not restore the ffERG photopic b-wave, showing the specificity of the RHO promoter.

Statistical ananlysis of the Visual acuity data shown in Figure 4.

| **Supplementry Table 1.** | |
|---|---|
| Two-way Repeated Measure ANOVA P values for Figure 4 | |
| Number of families | 6 |
| Number of comparisons per row family | 1 |
| Number of comparisons per column family | 6 |
| Alpha | 0.05 |
| Šídák's multiple comparisons test | Adjusted P Value |
| Photopic vs. Scotopic | |
| Wildtype | 0.416 |
| Untreated *Lrit3*$^{-/-}$ | 0.003 |
| AAV8 RHO::Lrit3 treated *Lrit3*$^{-/-}$ | 0.182 |
| AAV8 GNAT2::Lrit3 treated *Lrit3*$^{-/-}$ | 0.051 |
| Photopic comparisons | |
| Wildtype vs. Untreated *Lrit3*$^{-/-}$ | <0.001 |
| Wildtype vs. AAV8 RHO::Lrit3 treated Lrit3-/- | <0.001 |
| Wildtype vs.AAV8 GNAT2::Lrit3 treated Lrit3-/- | <0.001 |
| Untrd *Lrit3*$^{-/-}$ vs. AAV8 RHO::Lrit3 treated Lrit3-/- | >0.999 |
| Untrd *Lrit3*$^{-/-}$ vs.AAV8 GNAT2::Lrit3 treated Lrit3-/- | 0.023 |
| AAV8 RHO::Lrit3 treated Lrit3-/- vs. AAV8 GNAT2::Lrit3 treated Lrit3$^{-/-}$ | 0.02 |
| Scotopic | |
| Wildtype vs. Untrd *Lrit3*$^{-/-}$ | <0.001 |
| Wildtype vs. AAV8 RHO::Lrit3 treated Lrit3-/- | <0.001 |
| Wildtype vs.AAV8 GNAT2::Lrit3 treated Lrit3-/- | <0.001 |
| Untrd *Lrit3*$^{-/-}$ vs. AAV8 RHO::Lrit3 treated Lrit3-/- | 0.892 |
| Untrd *Lrit3*$^{-/-}$ vs. AAV8 GNAT2::Lrit3 treated Lrit3-/- | <0.001 |
| AAV8 RHO::Lrit3 treated Lrit3-/- vs. AAV8 GNAT2::Lrit3 treated Lrit3-/- | 0.032 |
